# Prokaryotic homologs of the major vault protein form engineerable vault nanocapsules

**DOI:** 10.64898/2026.11.04.764334

**Authors:** Januka S. Athukoralage, Peter H. Yoon, Amir Shanehsazzadeh, Eric Kauderer-Abrams, Nicholas T. Perry, Matthew G. Durrant

## Abstract

Vaults are the largest ribonucleoprotein particles in eukaryotic cells, built from 78 copies of the major vault protein (MVP), yet their function remains unknown four decades after their discovery. MVP homologs occur in prokaryotes and bacteriophages, but whether they form vaults is unknown. Here, we show that prokaryotic and bacteriophage MVP homologs assemble into vault particles. We surveyed approximately 15 billion proteins, found MVP homologs in 29 bacterial and 4 archaeal phyla, and used size-exclusion chromatography to rapidly screen and isolate vault particles of eight homologs expressed in *Escherichia coli*. Cryo-electron microscopy structures of the phage homolog L05 at high resolution reveal a shell of 96 MVP chains enclosing a volume ∼25% larger than human vaults, among the largest protein shells of defined size outside virus capsids. Prokaryotic MVPs typically carry seven wall repeats and eukaryotic MVPs nine, and adding wall repeats to L05 yields larger vaults. Prokaryotic vaults expand the known diversity of vault architecture and offer a tractable system for studying vault assembly and a size-tunable scaffold for cargo encapsulation.

## Introduction

Vaults are the largest ribonucleoprotein particles described in eukaryotic cells, roughly 1.8x larger than ribosomes^1,2^. First isolated from rat liver in 1986 (ref. 1), they have since been observed in other mammals, slime molds, and trypanosomes^1,3,4^, and a typical human cell is predicted to hold about 10,000 vaults^5^. Characterized mammalian vaults are built from 78 copies of the major vault protein (MVP), arranged as two half-shells that join at the waist^6–8^ to form a nanocapsule about 67 nm long and 40 nm wide, with a barrel-shaped wall and domed caps^4,6,9^. Each MVP chain comprises N-terminal repeat domains that stack to form the wall (wall repeats), a shoulder domain, a cap helix, and a C-terminal segment that completes the cap (Fig. 1a-b)^7,10,11^.

**Figure 1.**
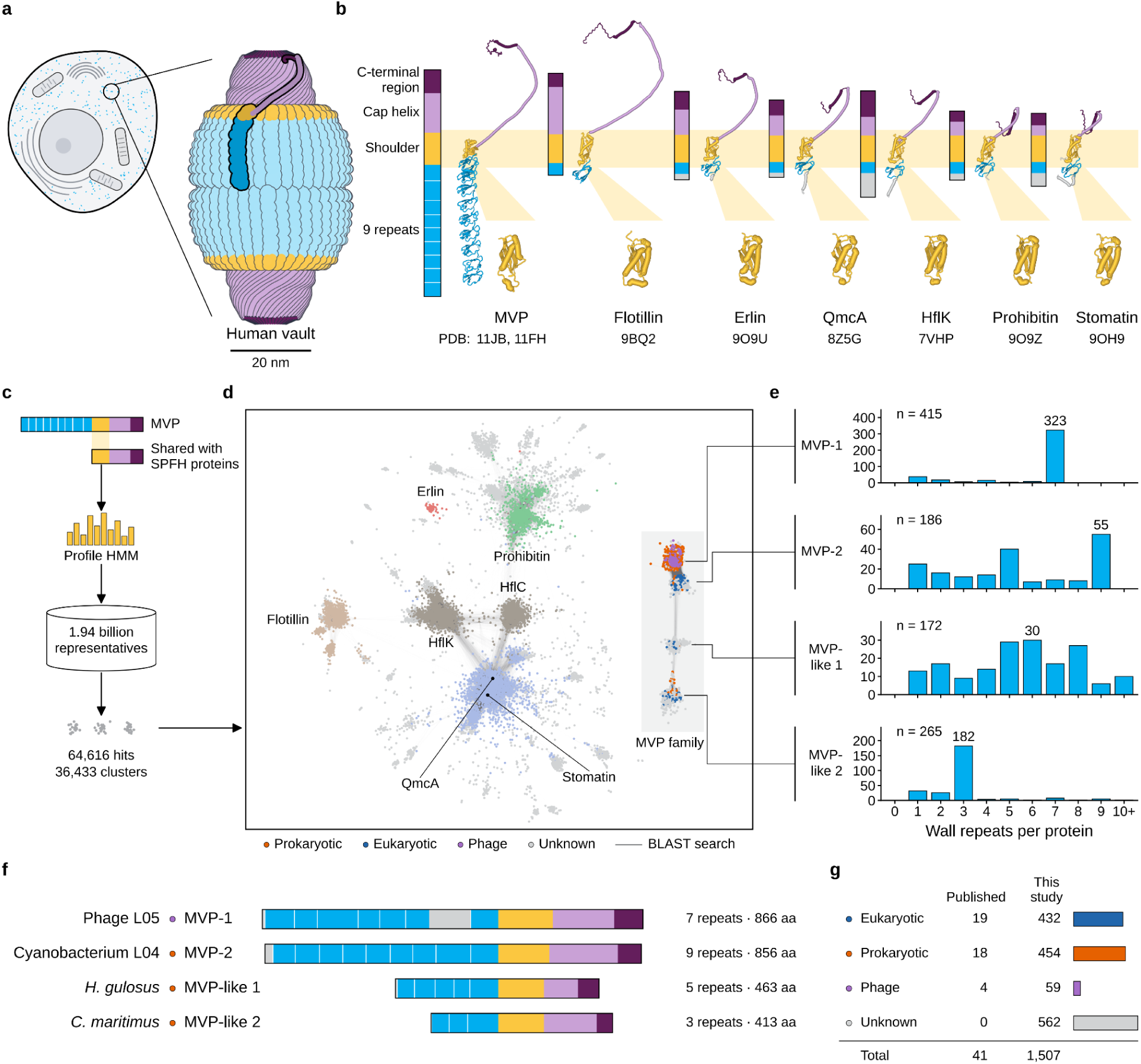
Major vault protein homologs occur across domains of life. (**a**) Schematic of a vault in a human cell. (**b**) Human MVP domain map compared to SPFH-domain-containing homologs. (**c**) Profile HMM search strategy for MVP homologs. (**d**) SPFH superfamily sequence similarity network. (**e**) Histogram of N-terminal repeat counts found within each MVP family cluster. (**f**) Representative domain architectures of MVP family members. (**g**) Number of new MVP homologs reported in this study.

In mammals, the shell encloses small vault RNAs (vtRNAs) and two associated proteins, PARP4 and TEP1 (refs. 3, 12). PARP4 is anchored inside the shell by its INT domain, which binds the wall at repeats 3 and 4 (ref. 13), and fusion to this domain is how cargo is loaded into engineered vaults^14–17^. Yet the function of vaults remains unknown: roles have been proposed in drug resistance, nuclear transport, signaling, and immunity^18–21^, but mice lacking individual vault components are viable^22–24^. Binding sites for NAD+ and ADP have been found inside the human vault^25^, and three recent studies report that a domain of TEP1 hydrolyzes NAD+^26–28^.

One route toward the function of vaults is to study their prokaryotic relatives. Studies of bacterial counterparts of human proteins, such as gasdermins, viperins, and cGAS–STING, demonstrated analogous function across domains of life^29–31^. In other cases, study of bacterial homologs led directly to the discovery of eukaryotic protein functions, such as TIR domains and sirtuins^32,33^. For vaults, uncovering the function of a prokaryotic homolog could shed light on eukaryotic vault function. MVP homologs have been identified in prokaryotes and bacteriophages^34–41^, and the bacterial homologs have been proposed to form vault-like assemblies^34^, but none has been shown to assemble into a vault.

Here, we survey the MVP family across approximately 15 billion proteins and show that prokaryotic and bacteriophage homologs assemble into vaults when expressed in *Escherichia coli (E. coli)*. The family spans 29 bacterial and 4 archaeal phyla but is rare, appearing in less than 1% of species-representative assemblies. Using size-exclusion chromatography (SEC) and negative stain electron microscopy (EM) as a rapid screen^42,43^, we found that seven homologs form vault-like particles and confirmed four as vaults by cryo-electron microscopy (cryo-EM). The structure of the phage homolog L05 reveals a shell consisting of 96 chains, 18 more than mammalian vaults. This assembly is shorter and wider than the human vault and encloses roughly 25% more volume. We identify contacts in the wall and cap that closed particles require, and show that adding wall repeats to L05 produces a nanocapsule that is 50% larger than the human vault. This work establishes prokaryotic vaults as a tractable system for studying vault assembly and function, and as a size-tunable scaffold for further engineering.

## Results

### Homologs of major vault protein occur across domains of life

MVP proteins include nine N-terminal repeats, a conserved central shoulder domain, and a C-terminal cap helix and cap disk (Fig. 1a)^7,10,11^. Past work focused on homology searches starting from full-length MVP sequences as queries and found a handful of prokaryotic MVP homologs. The shoulder domain, also known as the SPFH domain, is shared with a broader family of proteins called SPFH that form membrane-bound macromolecular complexes including stomatin, prohibitin, flotillin and HflK/C (Fig. 1b)^7,34,44–47^. To systematically map prokaryotic MVPs within this broader family, we constructed a profile hidden Markov model (pHMM) of the shoulder domain. We iteratively searched this against a previously compiled metagenomic database of approximately 15 billion protein sequences^48^ and clustered the resulting hits to identify novel MVP homologs (Fig. 1c-d; Methods)^49^.

All previously reported prokaryotic and eukaryotic MVPs clustered closely together in sequence similarity space, each respectively forming a distinct cluster we call MVP-1 and MVP-2 (Fig. 1d). Our clustering analysis also identified two more distantly related clusters that contain N-terminal repeats, a hallmark feature of MVP proteins. We denote these two clusters as MVP-like 1 and MVP-like 2. These two clusters were distributed across both prokaryotes and eukaryotes, but were largely of unknown host origin. We analyzed the number of N-terminal wall repeats in predicted AlphaFold2 structures for the proteins in these four clusters, and found that they varied considerably. MVP-1, where most prokaryotic MVPs lie, is defined by the presence of seven wall repeats and a long linker loop (∼80 aa) inserted between the sixth and seventh repeats (Fig. 1e,f). MVP-2, where canonical eukaryotic MVPs lie, has a more varied repeat number, most frequently nine, the same number as human MVP (Fig. 1e,f). MVP-like 1 and MVP-like 2 often retain membrane-anchoring features ancestral to the SPFH family, which may explain their distant relationship to the two MVP clusters (Supplementary Information (SI)). Altogether, this metagenomic survey increased the number of known MVP/MVP-like protein groups (50% identity) from 41 to 1,507, with the prokaryotic and eukaryotic MVP groups holding roughly equal numbers of proteins (Fig. 1g).

### MVP homologs from bacteria, phages, and archaea form vaults

Our survey showed that prokaryotic MVPs differ architecturally from eukaryotic MVPs, which raised the question of whether they form vault-like assemblies in a cellular context. To better understand the relationships within the broader MVP family, we built a phylogenetic tree from 489 representative sequences (SI; Methods). The clusters resolved into discrete phylogenetic groups spanning MVP-like genes found in bacteria, archaea, phages, and eukaryotes (Fig. 2a). To systematically survey these MVPs, we selected 44 proteins spanning 3 to 14 wall repeats from major MVP clusters and expressed all of them using cell-free systems and evaluated 25 heterologously expressed proteins for vault particle formation (SI; Methods). Since vault particles are so large (∼9 MDa), we opted to look for them in the excluded volume during size-exclusion chromatography, where large particles (>5 MDa) and aggregates are expected to elute (Fig. 2b; Methods). This method resembles earlier isolations of human vaults but reduces isolation time to only 90 minutes^42,43^.

**Figure 2.**
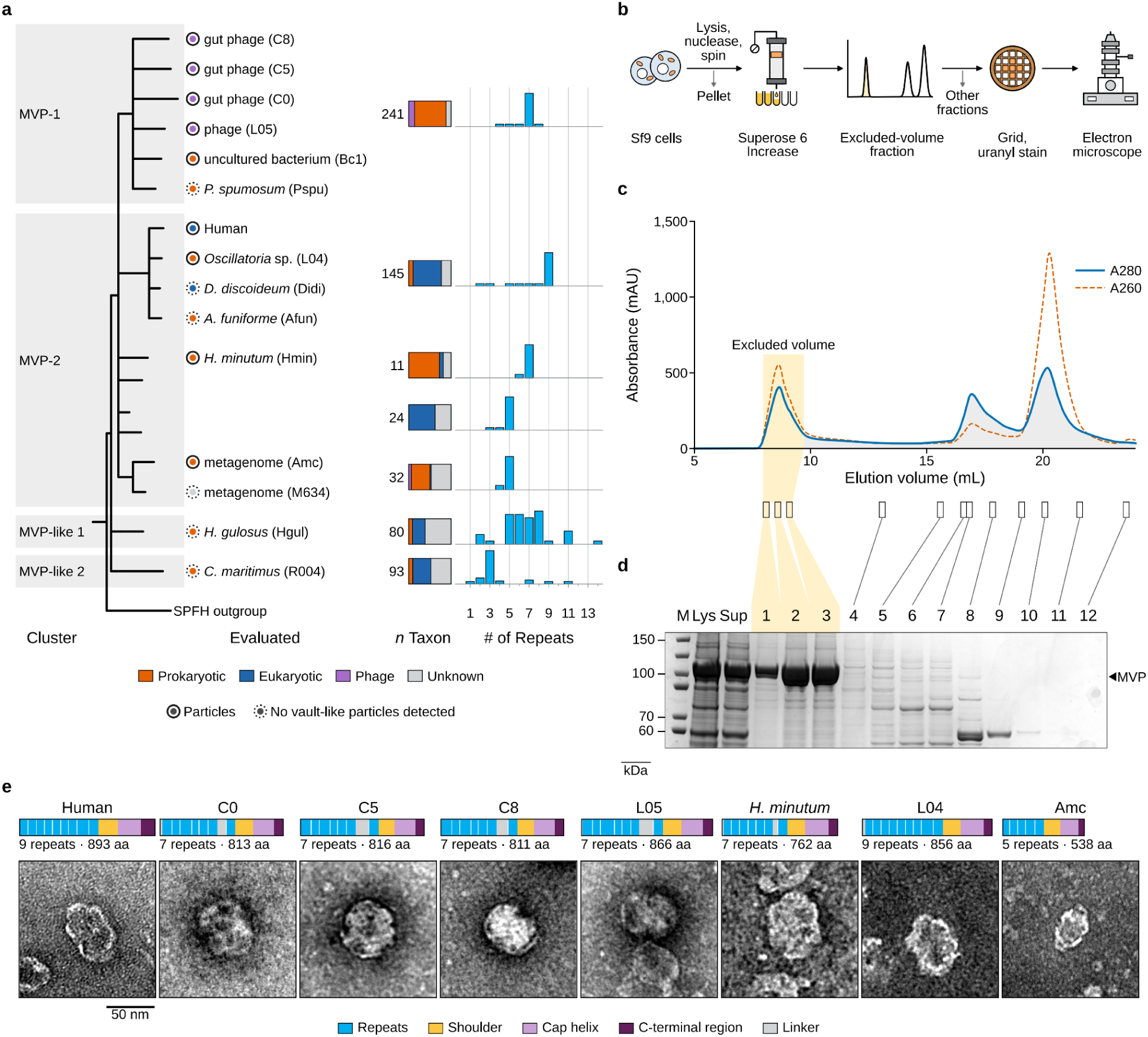
Excluded-volume capture of the human vault and MVP homologs made in *E. coli*. **(a)** Maximum-likelihood tree of MVP family proteins. **(b)** Vault particle isolation workflow. **(c)** Size-exclusion profile of human MVP purification. **(d)** SDS-PAGE of size-exclusion fractions. **(e)** Negative stain EM of excluded-volume fractions of human MVP and prokaryotic and phage MVP homologs.

To evaluate this method, we expressed human MVP in Sf9 cells and noted a discrete excluded-volume peak whose strongest band ran at the expected size on an SDS-PAGE gel (Fig. 2c-d). Negative stain EM of one fraction clearly showed intact human vault particles (Fig. 2e; SI). We next turned to the prokaryotic, bacteriophage, and other distant homologs of human MVPs, which we chose to express in *E. coli.* MVP homologs from four uncultured phages (C0, C5, C8 and L05), the myxobacterium *Hyalangium minutum* (Hmin), a cyanobacterium (L04), a probable archaeon (Amc), and eight other MVP candidates each gave excluded-volume material with a strong band near the expected size (SI). Negative stain EM of seven homologs showed vault-like particles, while the rest formed only rosettes (flower-like structures of opened vaults^50^), rings or aggregates indicative of half or malformed particles (Fig. 2e; SI). In contrast, an extract of cells carrying the empty vector, taken through the same procedure, gave a sparse grid with no barrels and no ring or arc-shaped fragments (SI). MVP homologs C0, C5, C8 and L05 gave compact particles, shorter than the human vault, whereas Hmin and Amc particles resembled the more elongated human vault in overall shape.

To better understand these vault-like particles, we used cryo-EM to evaluate C0, L05, Hmin, and Bc1. Cryo-EM micrographs confirmed that these particles closely resemble human vaults, with two halves joined at a waist and horizontal ridges along the body possibly corresponding to the wall repeats (SI). C0, L05 and Bc1 form more spherical structures, whereas Hmin forms lemon-shaped structures with tapered ends. Bc1 appeared to be ring-like by negative stain EM but yielded particles when imaged by cryo-EM, suggesting that negative stain alone cannot conclusively identify the presence of a vault particle. To our knowledge, these are the first defined particles formed by the assembly of MVP homologs from outside eukaryotes, and we refer to them as vault particles or vaults from this point on.

### A high-resolution structure of a bacteriophage-derived vault particle

To determine how a bacteriophage MVP homolog builds a vault, we solved the structure of the L05 particle by cryo-EM, obtaining a structure with a nominal resolution of 3.2 Å (SI; Methods). The L05 particle is composed of two halves that each contain 48 MVP chains that join at the waist to form a closed shell, in contrast to the 39 found in each half of a human vault (Fig. 3a-b). As a result, the L05 particle is wider than the human vault along the short axis (47.5 nm vs. 40 nm). Each chain follows the same order as human MVP, beginning with N-terminal repeats at the waist and terminating at the C-terminal cap. Notably, the linker loop between the sixth and seventh repeats conserved in the MVP-1 group was not resolved in our structure. Each L05 chain has seven wall repeats, two fewer than human MVP, which makes the L05 wall 7.2 nm shorter (28.1 nm vs. 35.3 nm; SI). Furthermore, the cap of L05 is flatter than that of the human vault (∼34° in L05, ∼52° in the human vault). Overall, these structural features give the L05 particle an egg-like shape that contrasts with the elongated lemon-like shape of the human vault (Fig. 3a-b).

**Figure 3.**
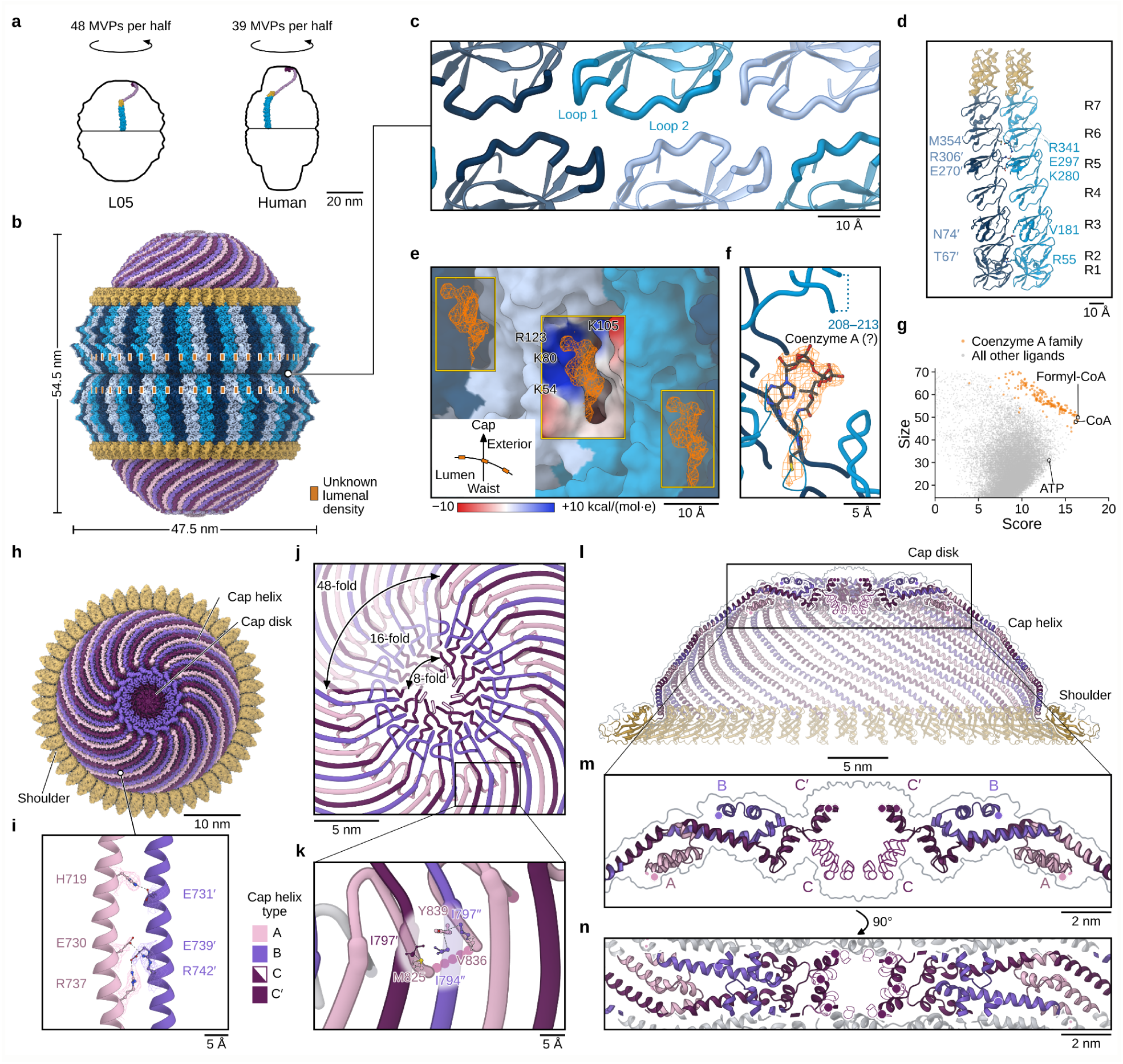
Cryo-EM reconstruction of the phage L05 vault particle. **(a)** Outlines of the L05 and human vaults (PDB 11JB). **(b)** Cryo-EM map of the closed L05 shell. **(c)** Vault waist interactions. **(d)** Wall seam formed by two MVP chains. **(e)** Wall pockets viewed from the lumen containing the unassigned density (orange). **(f)** Coenzyme A modeled into the wall pocket density. **(g)** Density fit score of ligands to unknown lumenal density. **(h)** Cap region of L05 vault. **(i)** Interaction between neighboring cap helices. **( j)** Symmetries of cap helix and cap disk. **(k)** Contacts at the cap disk. **(l)** Side view of the cap, depicting the shoulder, cap helices and cap disk. **(m)** Chain conformations A, B, C and C′ at the cap disk. **(n)** Top-down view of (l).

The two halves of the L05 vault meet at the waist through two loops of the first repeat, similar to the human vault (Fig. 3c)^6,25^. Interestingly, none of the residues lining this interface is shared between the two vaults, and the interface surface area of L05 is about half that of the human vault (∼225 Å² vs. ∼420 Å²). Above the waist, the repeats stack end to end within each chain, and neighboring chains lie side by side, meeting along a seam that runs the length of the wall (Fig. 3d). At this seam, the interface between neighboring chains covers about 2,300 Å² per chain and involves about 65 residues across all seven repeats. Across the seam, residues of one chain interact with residues of the neighboring chain. These include hydrogen bonding (R55–T67′), close packing (V181–N74′, M354′–R341) and salt bridges (K280–E270′, E297–R306′) (Fig. 3d; SI). Together, these contacts join the repeats of neighboring chains along the full height of the wall, closing the shell laterally.

Each repeat seam on the lumenal side of the wall carries a well-resolved density of uncertain origin not accounted for by the protein model, suggesting that the vault binds a small molecule (Fig. 3e-f). The density sits in a basic pocket between neighboring chains, formed by repeats 1 and 2. A computational search of 25,000 molecules against this density suggests that coenzyme A or one of its derivatives is the likely ligand (Fig. 3g; SI; Methods). Binding of nucleotide-like ligands has also been identified in the human vault, where NAD+ and related nucleotides bind the lumenal wall^25,26^. Each pocket is flanked by a short lumenal loop from repeat 3 that is unresolved in the model (residues 208–213). PARP4, the partner protein of the human vault, docks on the same lumenal face of the vault wall, at repeats 3 and 4 (refs. 25, 26). Together, these observations point to the lumenal face of the shell as a region where vaults may evolve the ability to bind small molecules and partner proteins.

The cap consists of two regions, the cap helix that begins to close the vault, and the cap disk at the poles (Fig. 3h). Neighboring cap helices pack side by side, with an interface of about 3,000 Å² that includes two salt bridges (R737–E739′, E730–R742′) and a hydrogen bond (H719–E731′) (Fig. 3i). Three different conformations of MVP subunits (A, B, C) arranged as trimers at the cap together make a 16-fold symmetrical cap disk (Fig. 3j). Notably, the C chains alternate between two forms (C and C′) at the center of the cap disk, further breaking the symmetry into 8-fold at the pole. The L05 cap is made of helices that narrow toward the pole in three steps: chain A folds back on itself into the lumen and tucks under adjacent chains B and C at about halfway from the pole (Fig. 3k). At the rim, the turned-back region of chain A packs against the ends of the next two cap helices (M825–I797′, V836–I797″, Y839–I794″ and I797″). Chain B extends further to the pole than chain A, but folds back on itself toward the exterior of the vault (Fig. 3l-m). The ends of chain C alone extend to the center of the cap disk (Fig. 3l-n). This tiered dome pattern resembles that of the mammalian vault, but notably differs in that it breaks the 48-fold symmetry twice and is composed purely of alpha helices rather than a central beta-barrel pore^10,11^. Together, this unique conformation of the cap suggests that several diverse cap-closing mechanisms have evolved from the same domain architecture and may be a hallmark feature distinguishing vault family members.

### Vault formation can be disrupted by single-residue substitutions

Neighboring MVP chains meet in three regions of the L05 shell: the wall, the shoulder and the cap. To test which of these contacts are required for assembly, we mutated each of these regions (Fig. 4a; SI; Methods). In the wall, we introduced or reversed charges along the seam. In the shoulder, we replaced both sides of an aromatic contact between neighboring chains with aspartate and reversed the charge of E654, which faces R592 of a neighboring chain. In the cap, we targeted the cap helix and the cap disk. We ran every variant on SEC and compared the relative amount of MVP eluted in the excluded volume, indicating complete particles or large aggregates, to other species migrating elsewhere. We imaged a subset of excluded fractions by negative stain EM to assess the presence of vaults (SI).

**Figure 4.**
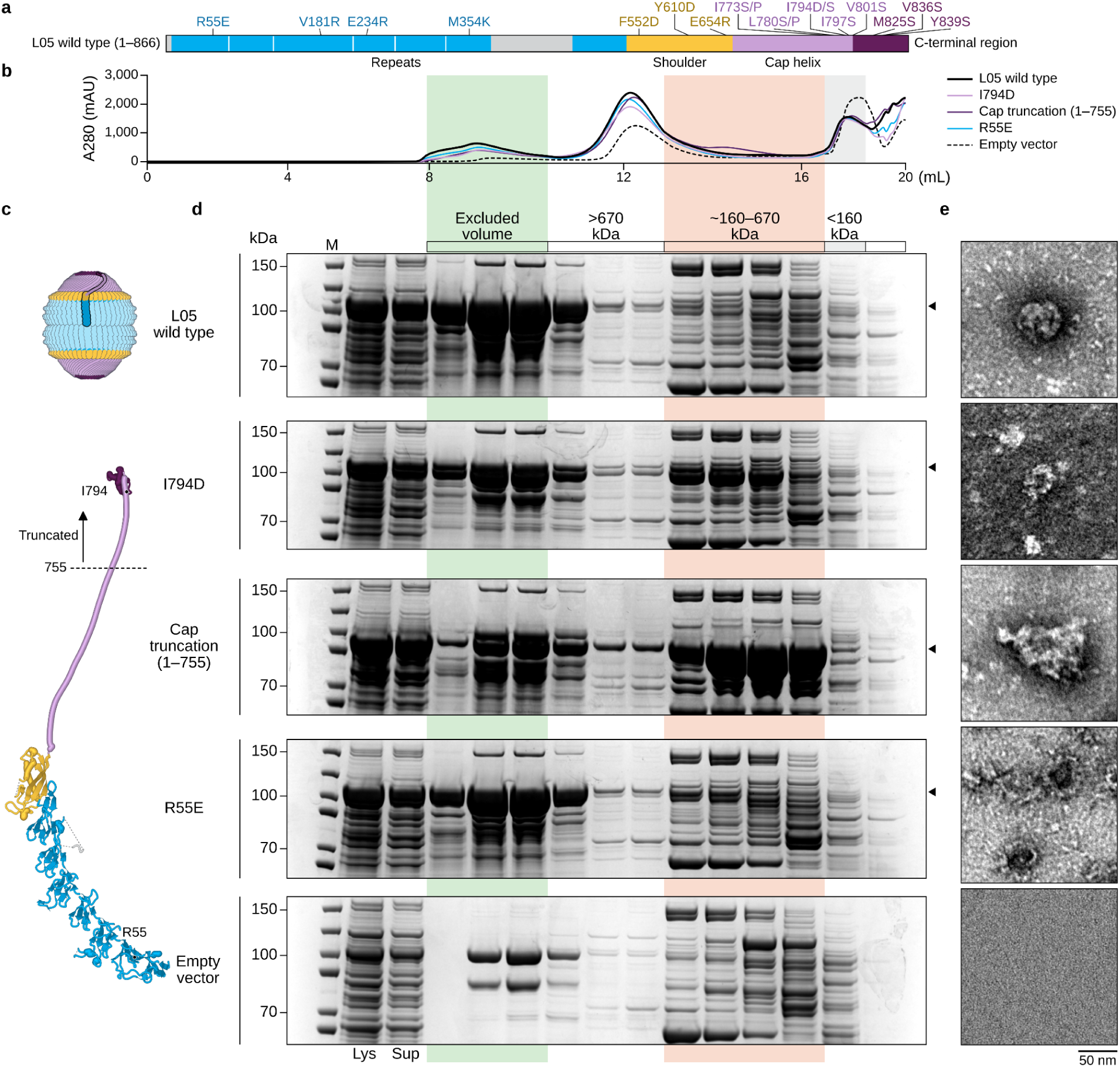
Site-directed mutations of L05 disrupt vault assembly. **(a)** Domain organization of the L05 phage MVP homolog with highlighted mutations. **(b)** Overlaid SEC traces of vault variant isolations. **(c)** L05 vault and predicted protein structure of a single MVP chain. **(d)** SDS-PAGE gel of different fractions of the SEC run. Lys, lysis; Sup, supernatant. **(e)** Negative stain EM of excluded-volume fractions in (d).

Substitutions in the wall or the cap disrupted closure, preventing complete vault assembly. All five substitution variants that we imaged still eluted in the excluded volume (Fig. 4b-d; SI), yet none formed vault-like particles (Fig. 4e; SI). Notably, a single charge reversal in the wall seam (R55E) or a single substitution in the cap helix (I794D) was enough to yield rosette-like particles instead of closed shells (Fig. 4d-e), as did multi-site substitutions in either region (SI). Likewise, substitutions at the cap rim produced large amorphous aggregates (SI).

Disrupting the cap helix also abolished vault assembly. The proportion of MVP in the excluded volume was greatly reduced for both substitution and truncation variants of the cap helix, and negative stain EM of such samples yielded amorphous aggregates rather than vaults (Fig. 4d; SI). Equivalent substitutions and truncations of the cap of a second MVP homolog, C0, similarly led to disruption of vault assembly (SI). In mammalian vaults, the cap helix mediates MVP self-association^51^ and stabilizes the assembled particle^7^. Our results suggest that this is a conserved feature among distantly related vault lineages, and that even single-residue mutations can dramatically impair assembly.

### Wall-repeat copy number is naturally variable and can tune vault size

MVP-family proteins vary considerably in their number of wall repeats (Fig. 1e). Mammalian MVPs carry nine repeats and most prokaryotic homologs carry seven, with some reaching up to fourteen (SI). We found that in some natural MVP sequences neighboring repeats are closely related at the primary sequence level (SI), suggesting that internal duplication can drive repeat expansion. For example, five bacterial and phage homologs with the same architecture as L05 carry several consecutive copies of repeat 4 that share up to 89% identity (Fig. 5a; SI).

**Figure 5.**
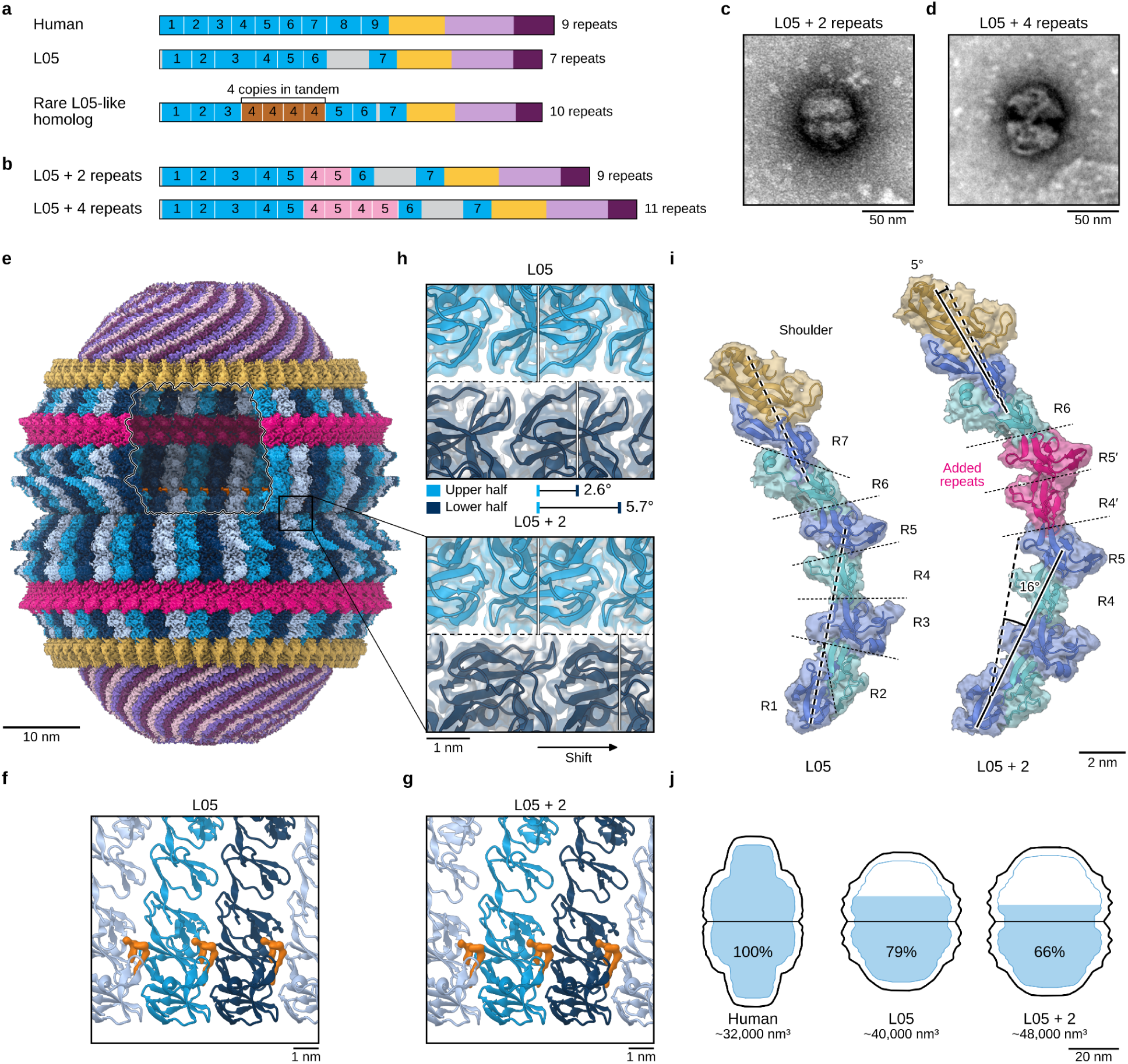
Natural and engineered expansion of the vault wall. **(a)** Domain organization of natural MVP homologs, with repeats and potential duplications indicated. **(b)** Domain organization of L05 variants with two and four added repeats. **(c-d)** Negative stain EM of (c) L05+2 and (d) L05+4 vaults. **(e)** Cryo-EM density of L05+2 vault. **(f-g)** Candidate small-molecule density in (f) L05 and (g) L05+2. **(h)** Comparison of L05 and L05+2 waist region. **(i)** Comparison of monomers of L05 and of L05+2. **( j)** Comparison of lumenal volume of vaults.

To recapitulate and test the effect of such duplications on vault assembly, we inserted extra copies of wall repeats 4 and 5 (residues 220 to 326) into L05 (Fig. 5b). L05 with two or four additional repeats (L05+2; L05+4) had similar SEC elution profiles to the wild-type vault, indicating that repeat expansion may not significantly perturb vault assembly (SI). When L05+2 and L05+4 were visualized by negative staining, we observed expanded vault particles with two lobes joined at the waist (Fig. 5c-d).

To see how the added repeats are accommodated, we determined the structure of L05+2 by cryo-EM to a nominal resolution of ∼3.5 Å (SI; Methods). L05+2 forms a closed shell that is longer and wider than L05 (58.5 by 49.4 nm vs. 54.5 by 47.5 nm) due to the additional repeats built into the wall (Fig. 5e). The wall pocket of L05+2 also holds the same unassigned density as that of L05 (Fig. 5f-g), further supporting that the L05 vault accommodates a small molecule such as coenzyme A. At the waist, the two halves are rotated by about 3° relative to L05 (Fig. 5h), suggesting that the added repeats change how the two halves meet. The additional repeats also change the curvature of the wall by 16° without disrupting the symmetry of the wall or the cap (Fig. 5i; SI). This results in a ∼20% increase in volume enclosed by the L05 vault, representing ∼50% more total volume than the human vault (Fig. 5j). Overall, these results show that the size, shape, and volume of vaults can be modulated by changing the number of their wall repeats, which makes repeat duplication an avenue for engineering vault nanocapsules, mirroring their natural evolution.

## Discussion

In this study, we show for the first time that prokaryotic and bacteriophage homologs of MVP assemble into vault-like structures. Earlier work has computationally identified some of these proteins in bacteria^34–36^, bacteriophages^37,38^, and archaea^39^ and predicted that they would assemble^34,35,39,40^. We confirm this prediction by cryo-EM, which shows closed particles of two joined halves for homologs from phages to bacteria. We further report cryo-EM structures of the phage-encoded L05 vault and an engineered variant. Overall, our work points to a broad and varied family of vaults in prokaryotes and their phages, with implications for what vaults do, how they are built, and how they can be engineered.

A clear biological function of human vaults has eluded scientists since their discovery, and we have likewise not yet established the function of prokaryotic vaults. However, finding vaults in all kingdoms of life, and being able to express and study them in bacterial cells, should make a function far easier to identify. In our survey, some subclades of MVPs lie in operons with partners involved in anti-phage defense (JAB/ubiquitin-like systems), RNA repair (RtcB), nucleotide metabolism (nucleosidases), and cyclic di-GMP signaling (SI). Our data further raise the strong possibility that phage vaults bind key metabolites like coenzyme A, analogous to NAD+ binding by human vaults. We propose that vaults may serve different functions in different lineages, perhaps united by a need to enclose their cargo. Further exploration of shared principles across the family by studying the MVPs reported in this work may shed light on the function of human vaults and their evolution.

Prokaryotic vaults conserve the broad architecture of eukaryotic vaults. In both, self-assembling monomers form two half-shells that join into the complete vault. Within this shared plan, chain number, repeat number and overall size vary. The L05 vault holds 96 chains rather than the 78 of mammalian vaults, each with fewer wall repeats, giving a shorter and wider shell with about 25% more lumenal volume. Adding two repeats to the L05 monomer increased the enclosed volume by a further 20% without disturbing assembly, and particles formed by other homologs, such as the lemon-shaped Hmin, suggest that more variations remain to be found. Structures of diverse vaults can therefore teach us the rules that set chain number, repeat number, curvature and cap form, opening a path to designer nanocapsules of defined size and shape.

Bioengineers have repurposed mammalian vaults to encapsulate various cargos with promising applications, such as enzymes^14^, small molecule drugs^15^, signaling molecules^13,52^, and cellular mRNA^53^. Similar to eukaryotic vaults that are engineered to encapsulate protein cargos fused to the PARP4 INT domain, prokaryotic vaults could be engineered in their repeat regions. Their repeat walls typically contain a long disordered loop that may be engineered to recruit binding partners. We show that diverse vaults can be produced, isolated and engineered rapidly in *E. coli,* which has proven to be difficult for mammalian vaults. Other delivery technologies, such as lipid nanoparticles and virus-like particles, became more useful as their size, shape and surface properties were explored and optimized^54,55^. Vaults begin as a closed shell of defined form, and the ability to alter that shell and to draw on the natural diversity of the family may likewise broaden their use for encapsulation and delivery.

## Acknowledgments

We thank Thomas Chi, Maxine Bi and Anna Howes of the Thermo Fisher Scientific Cryo-EM Drug Discovery Center for advice and for assistance with grid preparation and imaging. We thank Reena Zalpuri and Danielle Jorgens at the University of California Berkeley Electron Microscope Laboratory for advice and assistance with negative stain transmission electron microscopy. We thank Leonard Rome, Anqi Chen, Jonah Kallenbach, Raul Molina and Alex Lorenz for helpful discussions and comments on the manuscript. We thank FutureHouse, including Samuel Rodriques, Jessica Wu and Eli Moss, for supporting M.G.D. as a research fellow and for helpful discussions about vaults. We thank the life sciences team at Anthropic for their support. A portion of these data were produced by the US Department of Energy Joint Genome Institute (https://ror.org/04xm1d337; operated under Contract No. DE-AC02-05CH11231) in collaboration with the user community.

## Materials and Methods

### Sequence database and profile HMM search

MVP homologs were searched using a metagenomic database of ∼15 billion protein sequences^48^, represented by 1,939,242,578 cluster representatives at 50% identity (DIAMOND linclust^56,57^). A curated seed set of 512 sequences (489 of the MVP family and 23 SPFH proteins) was aligned with MAFFT 7.526 (ref. 58), trimmed to the shoulder region (residues 500 to 665 of *R. norvegicus* MVP). This alignment was further split into fourteen different alignments based on sequence phylogeny (one for all MVP sequences and one for each of nine MVP clades, and one for all non-MVP SPFH sequences and one for each of three SPFH clades), which were each turned into profile hidden Markov models (HMMs; pyhmmer 0.12.3 (ref. 59)). These HMMs were searched together with four SPFH models from Pfam 38.2 (ref. 60) (64,616 hits). A hit was counted as an MVP family member (i.e., MVP/MVP-like) if its best MVP profile scored at least 41.6 bits in a single domain and at least 10 bits above every SPFH model, and if the rest of the chain matched one of 18 additional custom HMMs that cover regions outside the shoulder domain (E ≤ 10⁻³), resulting in a total of 1,601 MVP family member candidate sequences.

### Sequence similarity network and cluster counts

A total of 64,540 MVP and SPFH profile hits and 1,303 reference sequences were pooled, and only sequences of at least 200 aa in length were kept for further analysis. The final set of 41,356 sequences was clustered with MMseqs2 18-8cc5c^61^ (mmseqs cluster --min-seq-id 0.5 -c 0.8 --cov-mode 0 --cluster-mode 0 -s 7.5), which resulted in 36,433 clusters. To perform clustering analysis, representative sequences from different clusters were selected. These included 1,429 sequences from all 1,427 clusters that contain MVP family sequences, 54 from clusters that contain reference SPFH sequences, and 6,000 from additional clusters drawn at random. The resulting 7,483 sequences were compared all against all with BLASTP^62^ (BLAST+ 2.17.0) and laid out with a custom force-directed program modeled on CLANS^49^. On this map the MVP family formed four clusters, which were named MVP-1, MVP-2, MVP-like 1, and MVP-like 2. To assess how many of the MVP/MVP-like sequences identified in the study were novel, these sequences were pooled with MVP sequences reported in the literature and grouped using MMseqs2 (mmseqs cluster --min-seq-id 0.5 -c 0.8 --cov-mode 0 --cluster-mode 0 -s 7.5; 1,945 clusters).

### Structure prediction and wall repeat counts

To analyze the number of wall repeats, the structures of 339 cluster representatives were predicted with AlphaFold2 (ref. 63) using ColabFold 1.6.1 (ref. 64). The nine repeats of PDB 4HL8 (ref. 8) were aligned to each structure with TM-align^65^ and with in-house code. Alignments with a TM-score^66^ of at least 0.45 were counted as repeats. For the 1,064 sequences without predicted structures, the number of repeats was inferred by sequence alignment to the closest of 293 full-length homologs with known counts.

### Phylogenetic analysis

Representative MVP/MVP-like and SPFH outgroup sequences were aligned with MAFFT L-INS-i^58^, and gappy columns were removed (gap threshold 0.5). The tree was inferred with IQ-TREE 3 (ref. 67) under LG4M^68^ with SH-aLRT^69^, aBayes^70^, and ultrafast bootstrap^71^ supports and rooted on the SPFH sequences.

### Prevalence and taxonomic distribution

In 111,922 GTDB species-representative assemblies^72^ (release 214.1 plus the species new in release 220; from OpenGenome2 (ref. 73)), every open reading frame of 100 codons or more was searched (pyhmmer 0.12.3) against three whole-chain HMMs. Genes that scored 100 bits or more and aligned from at least 120 profile columns before the shoulder were considered MVP family proteins, identifying 565 carriers in 26 bacterial and 3 archaeal phyla.

### Gene neighborhood analysis

Neighboring genes were analyzed at 3,507 loci (217 lineages at 50% approximate identity) of the MVP-1 cluster, each with a homolog of 750 or more residues and 10 kb or more assembled on both sides. Genes were predicted with pyrodigal-gv 0.3.2 (refs. 74–76) and annotated with hmmsearch^77^ against Pfam-A 38.2 at gathering thresholds. Loci were classified as phage-type or chromosome-type from geNomad marker genes (database v1.9). After deduplication with MMseqs2 (ref. 61), MVP neighboring genes were counted at 2,185 loci in 212 lineages (SI). Candidate operonic partners were curated manually on the basis of gene distance, conservation across lineages, and sequence diversity within groups.

### Prediction of whole-vault structures

For each of 22 proteins, whole vaults were modeled with 39, 45, and 48 chains per half (66 models; SI). Pairs of full-length chains were co-folded without templates with OpenFold3 0.5.0 (refs. 78, 79) and the OpenBind-0 weights80 (C8: OpenFold3 0.4.1 (ref. 78) and the preview-2 weights), through Anthropic’s inference kit81, from unpaired ColabFold alignments64. Predictions were retained if the chains lay side by side with continuous cap helices about 8 to 11.5 Å apart, and at least 10 were pooled per protein. Cα distances below 14 Å between neighboring chains found in at least half of the pooled predictions served as restraints to optimize one chain so that n exact copies about one axis satisfied them. The second half was generated by a two-fold axis, 1.5 Å from the first at the waist. Models were kept only if their handedness (tested at n = 39) agreed with the H. sapiens vault (PDB 11JB26), and only C0 was relaxed. For 13 proteins, the model at 48 used an earlier, smaller pool.

### Reference vault models and outlines

A predicted L05 model with 48 chains per half was built as described above (127 usable of 150 predictions; residues 8 to 798 of 866; not relaxed). Filtered to 20 Å, it provided the starting references for cryo-EM refinement of the L05 consensus and cap maps. For comparison with the modeled shells (SI), the H. sapiens vault was built from PDB 9BW6 (ref. 25) with its 39 assembly operators, and the L05 vault was the 96-chain cryo-EM model of this study. The C0 models (73 usable of 100 predictions) were refined in OpenMM 8.2 (ref. 82) with Amber ff14SB83 and OBC2 implicit solvent84. Outlines are the largest Cα radius in 0.25-nm slices along the particle axis, followed by the largest value over a running window of 1.75 nm.

### Constructs and cloning

Forty-four open reading frames encoding 43 distinct MVP homologs were codon-optimized for *E. coli* (SI). For cell-free expression, each was synthesized as linear double-stranded DNA carrying a T7 promoter, a ribosome-binding site and a T7 terminator (Twist Bioscience), and amplified by PCR. For expression in *E. coli*, untagged open reading frames were cloned into pET-21(+), and plasmids were verified by whole-plasmid sequencing. *H. sapiens* MVP was cloned into pFastBac1, and recombinant baculovirus was generated in Sf9 cells by standard procedures.

### Cell-free protein expression

Linear templates (10 ng/µL) were expressed in the PURExpress system (New England Biolabs) at 37 °C for 4 h. For fluorescent detection, reactions were supplemented with FluoroTect GreenLys tRNA (Promega)^85^. Reactions were separated by SDS-PAGE and imaged for in-gel fluorescence. Full-length bands were integrated with local background subtraction and normalized to a C0 MVP reference lane run on every gel. A construct was scored as full length when the full-length band was the strongest band in its lane (SI).

### Protein expression in insect cells and extract preparation

Sf9 cells in suspension at 1 × 10⁶ cells per mL were infected with 10% (v/v) passage-2 baculovirus and harvested by centrifugation after three days. Pellets were lysed for 15 min on ice in I-PER insect-cell protein extraction reagent (Thermo Fisher) containing EDTA-free protease inhibitors (Roche), at 5 µL per mg of wet cells. Lysates were centrifuged at 1,000 × g for 5 min, and the supernatant was treated with Benzonase (500 U/mL) for 20 min at room temperature. Extracts were then clarified at 20,000 × g for 20 min at 4 °C and passed through a 0.22 µm filter.

### Protein expression in *E. coli* and extract preparation

BL21(DE3) cells carrying pET-21(+) constructs were grown at 37 °C and 220 rpm in 100 mL of Terrific Broth containing carbenicillin (100 µg/mL), and expression was induced with 0.5 mM IPTG for 4 h. Proteins that were insoluble at 37 °C (Amc, R004, Bc1, Pspu, Didi and M634) were expressed at 16 °C. For these, cultures at an OD₆₀₀ of 0.6 were chilled in ice water for 30 min, induced with 0.5 mM IPTG, and shaken at 16 °C and 220 rpm overnight. Cells were harvested by centrifugation at 4,000 × g for 15 min at 4 °C and stored at −80 °C. Pellets were suspended at 3 µL per mg of wet cells in B-PER Complete (Thermo Fisher) supplemented with 150 mM NaCl, 1.5 mM MgCl₂, EDTA-free protease inhibitors (Roche) and Benzonase (Sigma-Aldrich), and incubated for 20 min at room temperature. Lysates were clarified at 21,000 × g for 20 min at 4 °C and passed through a 0.22 µm polyethersulfone filter. L05+2 and L05+4 were not filtered in this manner.

### Size-exclusion chromatography

Clarified extracts (500 µL) were applied to a Superose 6 Increase 10/300 GL column (Cytiva) on an ÄKTA pure 25 (Cytiva) at 4 °C and 0.75 mL/min, in buffer containing 50 mM Tris-HCl pH 7.5, 150 mM NaCl and 1 mM MgCl₂. Absorbance was recorded at 280 and 260 nm, and 0.25 mL fractions were collected. The column was calibrated with Blue Dextran 2000 (10 mg/mL) and eight protein standards (Gel Filtration Calibration Kits LMW and HMW, Cytiva) under the same conditions. Blue Dextran eluted at 8.96 mL, thyroglobulin (669 kDa) at 13.61 mL, ferritin (440 kDa) at 15.43 mL and aldolase (158 kDa) at 16.7 mL. Sizes marked on chromatograms are apparent sizes relative to these standards. The first peak of each extract was defined as the excluded volume, corresponding to fractions eluting between 7.97 and 9.72 mL.

### SDS-PAGE and in-gel fluorescence

Proteins were separated on NuPAGE 4–12% Bis-Tris gels (Thermo Fisher) in MOPS buffer at 200 V for about 50 min. Samples were mixed with NuPAGE LDS sample buffer and reducing agent (Thermo Fisher) and heated before loading, at 70 °C for 10 min. Lysates and clarified extracts were diluted 1:10 (Sf9) or 1:20 (*E. coli*) before loading, and fractions from the most concentrated regions of the chromatogram were diluted four-fold where indicated. PageRuler Unstained Protein Ladder (Thermo Fisher) was used as the size marker on all gels. For in-gel fluorescence, BenchMark Fluorescent Protein Standard (Thermo Fisher) was loaded in the same lane. For cell-free reactions labeled with FluoroTect GreenLys, 2 µL of each reaction was loaded, and gels were imaged wet, directly after electrophoresis and before staining, for BODIPY-FL fluorescence on a ChemiDoc MP (Bio-Rad) using the fluorescein channel with a 60 s exposure. Gels were then stained with Coomassie and imaged on the same instrument. All images were recorded as 16-bit files. Uncropped scans are provided in the SI.

### Negative-stain electron microscopy

Excluded-volume fractions were diluted 20to 50-fold in column buffer and applied to grids without concentration. Formvar-carbon coated 300-mesh copper grids (01753-F, Ted Pella) were glow-discharged in air at 15 mA for 10 s (PELCO easiGlow, Ted Pella) directly before use. Sample (5 µL) was adsorbed for 1 min and wicked off with Whatman no. 1 filter paper. Grids were washed on three drops of double-distilled water, stained with 5 µL of 2% (w/v) aqueous uranyl acetate for 1 min, wicked dry and air-dried for 2 to 4 min. Grids were imaged on a Tecnai 12 transmission electron microscope (FEI) at 120 kV with a Rio16 CMOS camera and Gatan Microscopy Suite software (Gatan). Between 2 and 10 fields were recorded per sample. Micrographs (4,096 × 4,096 pixels) were calibrated from the scale bar recorded with each frame. Excluded-volume fractions from cells carrying the empty vector were prepared and imaged in the same way.

### Cryo-EM sample preparation and data collection

Excluded-volume fractions of L05, L05+2, Hmin, C0, and Bc1 were pooled and concentrated at 4 °C in 100-kDa centrifugal concentrators (Amicon Ultra, Millipore). Samples (2.5 µL) were applied to glow-discharged Quantifoil R 1.2/1.3 300-mesh grids (gold for L05; copper for L05+2 and Hmin). Grids were blotted in a Vitrobot Mark IV (Thermo Fisher Scientific; 5 °C, 100% humidity, blot force 0, 3 s) and plunge-frozen in liquid ethane. Movies were collected with EPU at the Thermo Fisher Cryo-EM Drug Discovery Center on a Krios G4 microscope at 300 kV with a Falcon 4i detector and an energy filter (10-eV slit). L05 was imaged at 130,000× (3,041 movies; 0.937 Å per pixel; 54.3 e⁻/Å²; defocus −0.5 to −1.5 µm), L05+2 at 130,000× (16,824 movies; 0.937 Å per pixel; 53.5 e⁻/Å²; defocus −0.8 to −2.0 µm), Hmin at 130,000× (8,080 movies; 0.937 Å per pixel; 53.0 e⁻/Å²; defocus −0.5 to −1.5 µm), Bc1 at 130,000× (201 movies; 0.937 Å per pixel; 53.5 e⁻/Å²; defocus −0.8 to −2.0 µm), and C0 at 81,000× (4,091 movies; 1.6 Å per pixel; 20 e⁻/Å²). Individual particles (SI) were cropped from micrographs that had been motion-corrected either in cryoSPARC Live v5.0.7 (L05, Hmin, and C0) or in RELION (Bc1).

### Image processing and reconstruction

For L05, 3,041 movies were motion-corrected with the RELION 5.0.1 (refs. 86, 87) implementation of MotionCor2 (refs. 88, 89), and contrast transfer function (CTF) parameters were fit with CTFFIND 4.1.14 (ref. 90). Topaz^91^ trained from manually selected particles was used to pick 34,337 particles from the 3,007 micrographs that were CTF fit to 6 Å or better. 2D classification retained 28,358 particles, and 19,389 of them were complete vaults. The complete vaults were used for three-class ab initio reconstruction without symmetry (3.75 or 1.87 Å per pixel). Four-class 3D classification of the 28,358 particles without symmetry, with references that were derived from ab initio maps, yielded 11,945 vaults that were closed and also complete in 2D classification. Refinements of entire vaults without symmetry from a featureless reference preferentially showed 48 repeats (ribs) around the wall. Sorting of signal-subtracted caps among the 48 positions showed a 16-fold cap, and refinements in which this symmetry was relaxed^92,93^ showed an 8-fold tip.

Ab initio maps did not distinguish the vault from its mirror image, so refinement was started from a 20-Å smoothened shell of 48 copies per half of a predicted model (see Reference vault models and outlines). The hand was confirmed by fitting ideal α-helices and their mirror images. After refinement under D16, CTF refinement^94^, and 3D classification-based particle clean-up with decoy references, 11,675 vaults yielded a consensus map at 3.50 Å (SI). Overlapping regions were then extracted by symmetry expansion and localized reconstruction^95,96^ and refined locally without symmetry. Local refinement reached 2.95 Å (waist), 3.02 Å (barrel), 3.39 Å (shoulder), and 3.24 Å (cap helices), and 23,404 caps, from 11,702 separately selected vaults, reached 3.30 Å under C16. The nominal resolution of 3.2 Å is the mean of these five values. The tip was reconstructed under C8, and the maps were merged into a composite map.

For L05+2, 16,824 movies were motion-corrected and CTF parameters were fit as for L05. Template picking yielded 114,327 particles from the 16,781 micrographs with a good CTF fit (6 Å or better). An ab initio model was generated without symmetry from 617 particles of closed-vault 2D classes and provided the initial reference. Particles were then sorted by 3D classification without symmetry (four classes), and the class containing closed vaults was kept (17,779 vaults). Refinement started from a 20-Å reference that was computed from the lengthened L05 model, which supplied the 48 ribs, the hand, and the outline. Refinement of whole vaults under imposed D16 symmetry did not reach the Nyquist limit (8.1 Å at best, from pixels binned to 3.75 Å) and was not pursued further (SI). In contrast, when each half was treated as a separate particle, the 35,558 halves reached 4.2 Å under C48 at 1.25 Å per pixel. Three overlapping regions of the wall were extracted from the halves after 48-fold expansion, refined locally without symmetry, and merged into a composite map. Local refinement to convergence of the sub-particles that moved as their neighbors did reached 3.33 Å (barrel), 3.58 Å (middle), 3.45 Å (shoulder and cap), and 3.8 Å (waist). The 12,446 caps that were placed with the highest confidence reached approximately 3.5 Å under C16 within 100 Å of the axis. The entire vault was generated from one half and its copy about a two-fold axis. The relative rotation of the two halves and the plane in which they meet could be determined because the map of the halves also contained density for the partner half. Refinement to convergence increased the difference between the half maps in the shells from 4.7 to 3.5 Å (1.09to 1.89-fold in the wall), indicating some fitted noise (SI).

All refinements followed the gold-standard procedure, in which the particles of each data set were divided into two half-sets that were refined and reconstructed separately^97^. Resolutions of the reported maps correspond to a Fourier shell correlation (FSC) of 0.143 between the two resulting half maps after mask correction^98^. Steps for which RELION has no job type were performed with custom Python scripts (SI). These steps included sub-particle extraction, cap sorting, geometric mask construction, composite map assembly, the check for fitted noise, and directional FSC calculation.

### Model building, refinement, and validation

For L05, an AlphaFold2 model (ColabFold 1.6.1) was docked into the map, the cap helix was built from ideal helices, and the end of the chain was built in the C16 and C8 maps. Model building was performed iteratively with rebuilding in Coot 1.3.3 (ref. 99), refinement in Servalcat 0.4.131 (refs. 100, 101) under strict symmetry, scoring, and residue-by-residue inspection. D8 symmetry generates the 96 chains of the model from six, which adopt four conformations (A, B, C, and C′) that differ beyond residue 788. Each chain contains 717 to 724 of the 866 residues. The final refinement was run against earlier maps from longer local refinements. To model L05+2, the L05 chain was split between residues 326 and 327, and a second copy of residues 220 to 326 was added (residues 327 to 433 of L05+2; residues 327 to 866 of L05 became 434 to 973). The pieces were docked into the map as rigid units, joined, and refined with Servalcat (wall to 3.5 Å; cap to 3.7 Å).

### Use of AI tools

Analyses were carried out with various Claude (Anthropic) models across several interfaces and harnesses, including Claude Science, Claude Code, Claude Desktop, and other internal tooling. Claude wrote all code and carried out all computational analyses in this study. The authors directed the analysis and take responsibility for the content.

